# Impact of cell size on morphogen gradient precision

**DOI:** 10.1101/2022.02.02.478800

**Authors:** Jan A. Adelmann, Roman Vetter, Dagmar Iber

## Abstract

Tissue patterning during embryonic development is remarkably precise. We numerically determine the impact of the cell diameter, gradient length, and the morphogen source on the variability of morphogen gradients and show that the positional error increases with the gradient length relative to the size of the morphogen source, and with the square root of the cell diameter and the readout position. We provide theoretical explanations for these relationships, and show that they enable high patterning precision over developmental time for readouts that scale with expanding tissue domains, as observed in the *Drosophila* wing disc. Our analysis suggests that epithelial tissues generally achieve higher patterning precision with small cross-sectional cell areas. An extensive survey of measured apical cell areas shows that they are indeed small in developing tissues that are patterned by morphogen gradients. Enhanced precision may thus have led to the emergence of pseudostratification in epithelia, a phenomenon for which the evolutionary benefit had so far remained elusive.

## Introduction

During embryogenesis, cells must coordinate complex differentiation programs within expanding tissues. According to the French flag model [1], morphogen gradients define pattern boundaries in the developing tissue based on concentration thresholds. Exponential functions of the form

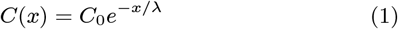

approximate the shape of measured morphogen gradients very well [2–9]. For such gradients, the mean readout position

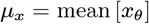

and the positional error

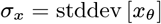

of the domain boundary positions

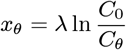

in different embryos depend on the variation in the decay length *λ* and in the amplitude *C*_0_ relative to the concentration threshold *C_θ_*. Strikingly, the positional error of measured morphogen gradients has been reported to exceed that of their readouts [10, 3, 11]. Several theories have been proposed to explain the high readout precision despite inevitable noise and variation in morphogen gradients and their readout processes. They include temporal and spatial averaging, self-enhanced morphogen turnover, the use of opposing gradients, dynamic readouts, and cell-cell signalling [10, 3, 12–15, 11, 16–20]. In zebrafish, where cells are rather motile, cell sorting and competition can further enhance boundary precision [21–23]. Here, we study patterning precision conveyed by morphogen gradients in epithelia and leave the effect of precision-enhancing processes in the morphogen readout for future work.

**Figure 1:**
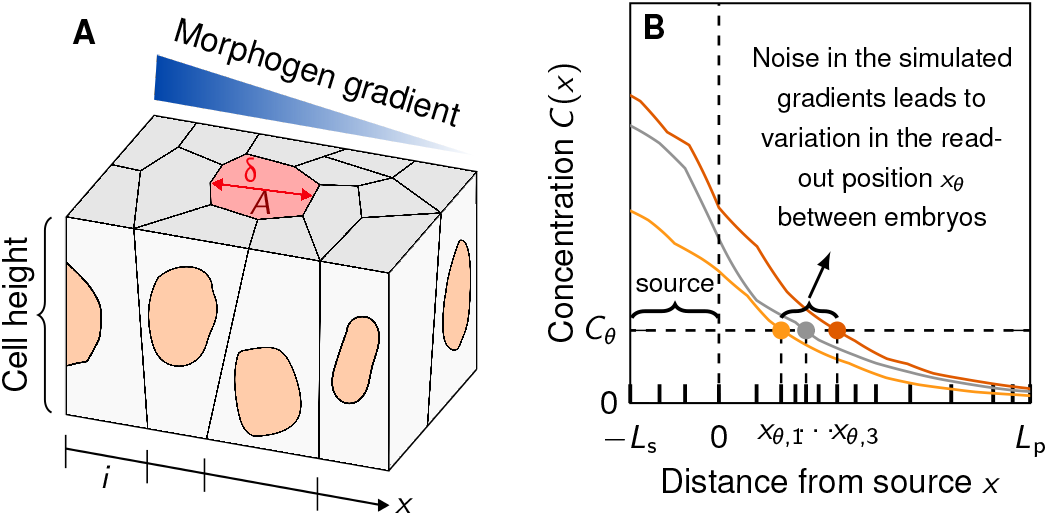
Patterning in epithelial tissues with variability in the morphogen kinetics and cell size. **A** Schematic of an epithelial layer of cells (index *i*) with cross-sectional area *A* and diameter *δ* along the patterning axis *x*. **B** Schematic of positional variability resulting from the readout of noisy gradients in a cellular domain, split into a morphogen-secreting source of length *L_s_* and a patterning domain of length *L*_p_.

A recently developed numerical framework estimates how much variability in and between morphogen gradients can be accounted for by cell-to-cell variability reported for morphogen production, decay, and diffusion [24]. In this article, we extend the model to take a different perspective on the precision of gradient-based patterning in cellular tissues. We analyse the impact of various length scales present in the epithelium, such as the cell diameter and source size, as well as spatial averaging, on morphogen gradient variability, finding that positional accuracy is higher, the narrower the cells and the larger the morphogen source.

We approximate the patterning axis by a discrete line consisting of two subdomains, a source domain on the interval –*L*_s_ ≤ *x* ≤ 0 and a patterning domain on the interval 0 ≤ *x* ≤ *L*_p_, each divided into sub-intervals *i* representing individual epithelial cells with diameter *δ_i_* in 1D, or cross-sectional areas *A_i_* in 2D (Fig. 1A). Noisy exponential gradients were generated by numerically solving the one-dimensional steady-state reaction-diffusion boundary value problem [24]

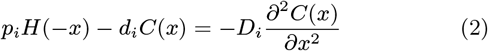

with zero-flux boundary conditions

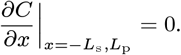

**Figure 2:**
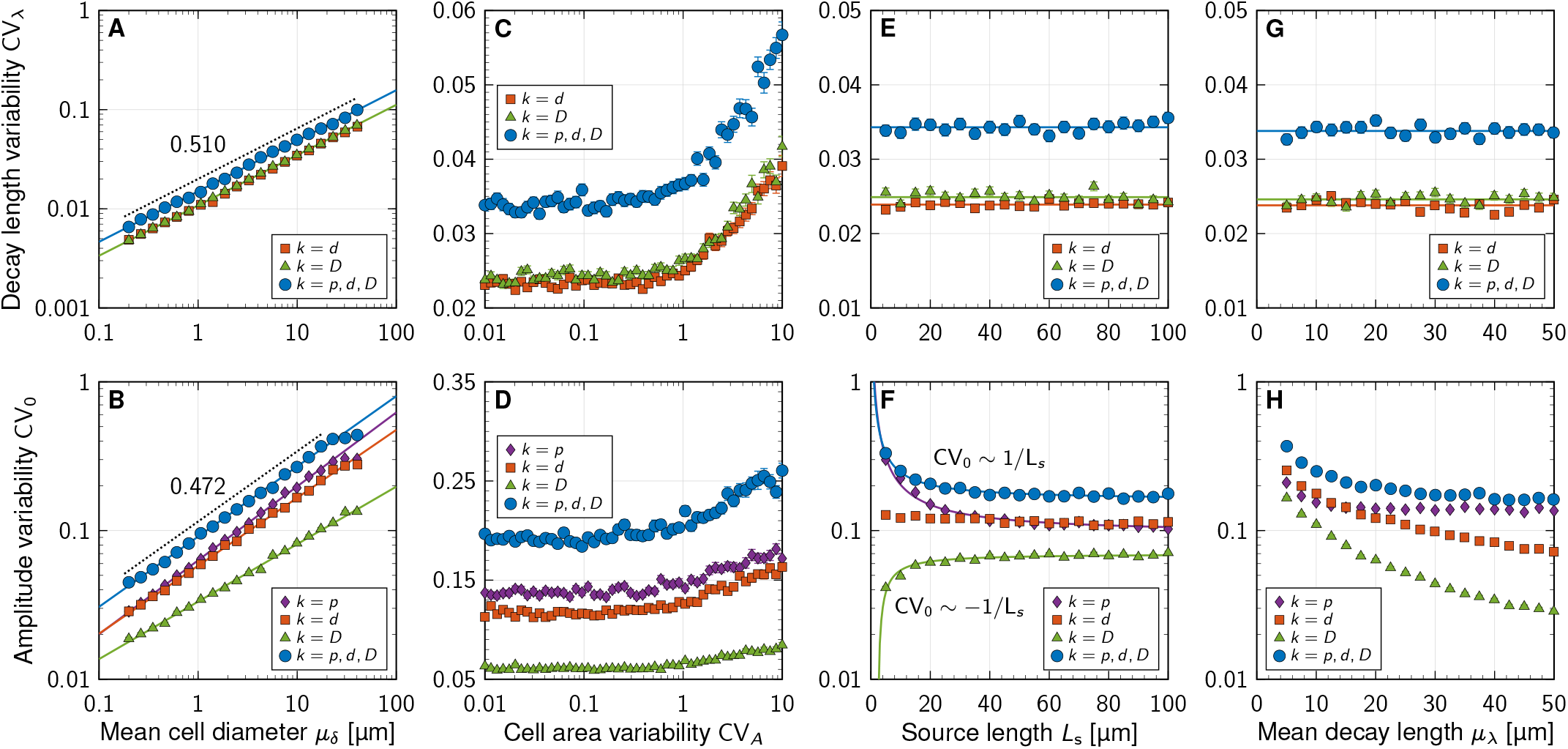
Impact of cell size, source length and gradient length on morphogen gradient variability. **A,B** Scaling of gradient variability with the cell diameter at fixed kinetic variability *CV_p,d,D_* and fixed cell area variability CV_*A*_. Fitted power-law exponents are indicated. **C,D** Gradient variability is largely unaffected by cell area variability as long as CV_*A*_ < 1 and increases only for greater values. **E, G** Variability in the gradient decay length is unaffected by the morphogen source size and the mean gradient decay length. **F** Except when only the diffusion coefficient varies (green), greater source lengths reduce morphogen amplitude variability. **H** Gradients with greater decay length contain less amplitude variability. Each data point in **A**–**H** corresponds to the mean ± SEM of *n* = 10^3^ independent simulations, with kinetic variability only in the parameters indicated by different symbols: CV_*k*_ = 0.3, CV_¬*k*_ = 0. Cell area variability: CV_*A*_ = 0.5 except in **C,D**. Domain sizes: *L*_s_ = 25 μm except in **E, F**, *L*_p_ = 250 μm except in **G, F**. The effect of CV_*p*_ on CV_λ_ is minuscule, 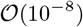, and therefore not plotted in the top row. See supplementary Table S1 for fit parameters.

Eq. 2 contains a source with production rates *p_i_*, a linear sink with degradation rates *d_i_*, and models morphogen transport by Fickian diffusion with effective coefficients *D_i_* subscripts *i* indicate that they vary from cell to cell. The Heaviside step function *H*(–*x*) ensures that morphogen production only occurs in the source, whereas degradation is assumed to take place over the whole domain. The kinetic parameters *k* = *p,d,D* were drawn for each cell independently from log-normal distributions. This assumes statistical independence of neighbouring cells; we will later relax this assumption by introducing spatial correlation. The distributions had prescribed mean values *μ_k_* and respective coefficients of variation CV_*k*_ = *σ_k_/μ_k_* analogous to [24]. We fixed molecular variability at the physiological value CV_*k*_ = 0.3 [24] here.

As a new source of noise, we introduced cell size variability. Since the cell area distributions in the *Drosophila* larval and prepupal wing discs, and in the mouse neural tube resemble lognormal distributions [25, 26], we drew individual cell areas *A_i_* independently from a log-normal distribution with prescribed mean *μ_A_* and coefficient of variation CV_*A*_. This allowed us to evaluate the impact of cell-to-cell variability in the production, degradation and diffusion rates *p_i_*, *d_i_, D_i_*, as well as in the cell cross-sectional areas *A_i_*, on gradient variability (Fig. 1B).

## Results

### Gradient variability increases with cell size, but not with physiological levels of cell area variability

We quantify relative variability or uncertainty of a positive quantity *X* by its coefficient of variation CV*v* = *σ_X_/μ_X_*, where *μ_X_* and *σ_X_* denote the mean and standard deviation of *X*, respectively. For the local morphogen concentration, this is CV_*C*_. Alternatively, one can fit Eq. 1 to each generated morphogen gradient (see Methods) and quantify CV_λ_ and CV_0_ of the two free parameters *λ* and *C*_0_ individually.

Our simulations reveal that an increase in the average cell diameter *μ_δ_* leads to greater variability in *λ* and *C*_0_ (Fig. 2A,B), according to power laws

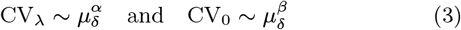

with exponents *α* = 0.510 ± 0.004 (SE, Fig. 2A, blue curve) and *β* = 0.472 ± 0.005 (Fig. 2B, blue curve). The amplitude variability CV_0_ plateaus when *μ_δ_* ≥ *L*_s_, because the source defaults back to a single cell in this case. Square-root scaling for the decay length variability (*α* =1/2) follows theoretically from the law of large numbers and is consistent with the inverse-square-root scaling reported for the dependency of CV_λ_ on the patterning domain length *L*_p_ at fixed cell size [24]. Together, this suggests that

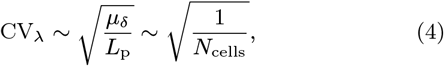

where *N*_cells_ is the (mean) number of cells along the patterning axis. Similarly, morphogen sources composed of more, smaller cells buffer cell-to-cell variability in morphogen kinetics more effectively, leading to the observed reduction in amplitude variability CV_0_. Smaller cell diameters thus lead to smaller effective morphogen gradient variability.

**Figure 3:**
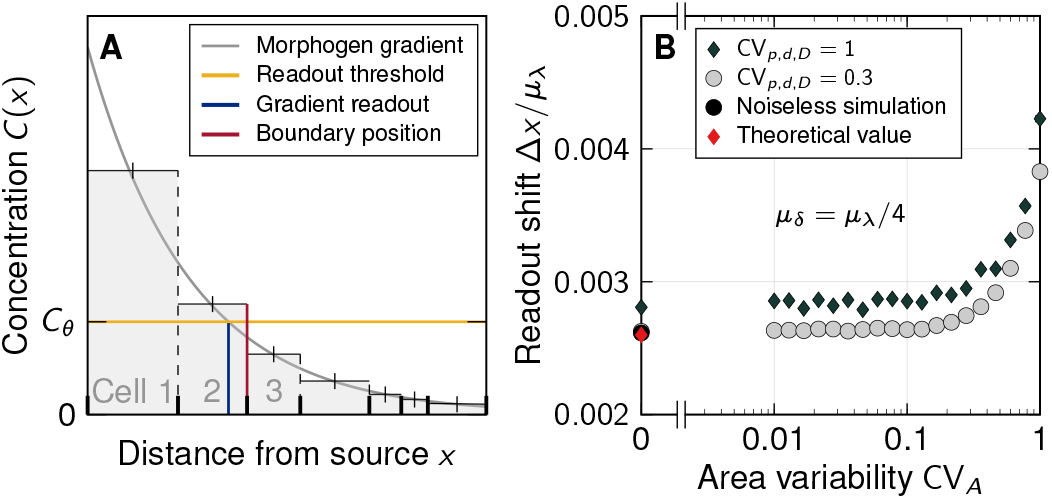
Readout position of exponential gradients is barely shifted by spatial averaging. **A** Cell-based readout of a morphogen gradient. A concentration threshold *C_θ_* (yellow) defines a readout position *x_θ_* (blue). If cells read out cell-area-averaged concentrations, the effectively sensed concentration profile is a step function (grey). Pattern boundaries form at cell edges (red). For illustrative purposes, the cell size is exaggerated compared to the gradient decay length. **B** Cell-area-averaged readout of exponential gradients results in a small shift Δ*x* compared to readout at the cell centroid.

Cell-to-cell variability in the cross-sectional cell area *A* does not affect the gradient variability as long as CV_*A*_ < 1 (Fig. 2C,D). Only for extreme cell area variability exceeding 1, the variability in *λ* grows (Fig. 2C). However, we are not aware of any reported CV_*A*_ > 1 [26–29]. Consequently, cell size has a considerable impact on gradient variability, while physiological levels of variability in the cell area do not contribute to gradient imprecision.

A larger source or gradient length reduces only the amplitude variability, but does not affect the decay length variability (Fig. 2E–H). Amplitude and gradient decay length variability is reduced in a source that is composed of many cells with a small mean diameter (see Supplementary Material, Fig. S5). The parameter values in the simulations correspond to those reported for the mouse neural tube (*μ_λ_* = 20 μm, *μ_δ_* = 5 μm, *L*_s_ = 5*μ_δ_*, *L*_p_ = 50*μ_δ_*). At these values, source sizes above 25 μm and gradient decay lengths above 20 μm barely reduce amplitude variability. Sonic hedgehog (SHH) in the neural tube is secreted from both the notochord and the floor plate, while Bone morphogenetic protein (BMP) is secreted from both the ectoderm and the roof plate. Intriguingly, while the SHH-secreting notochord shrinks over time, it still measures about 30 μm in width by the 5 somite stage [30], and the SHH-secreting floor plate then emerges in the ventral part of the neural tube and widens over time [31]. The gradient length remains constant at about *μ_λ_* = 20 μm [8, 11], the largest value for which the positional error remains small at a large distance (12*μ_λ_* = 240 μm) from the source. The source size thus assumes the smallest and the gradient decay length the largest value for which morphogen gradient variability remains small.

### Readout position is barely shifted by spatial averaging

Since cells can assume only a single fate, domain boundaries must follow cell boundaries (Fig. 3A). We sought to quantify the impact on the readout position if epithelial cells average the signal over their entire apical cell surface. Assuming that cells have no orientational bias, we can approximate cell surfaces as disks with radius *r* = *μ_δ_*/2 about a centre point *x*_0_. If threshold-based readout operates on the averaged concentration, the effective readout domain boundary is shifted along the exponential concentration gradient to *x*_0_ = *x_θ_* + Δ*x* by the distance

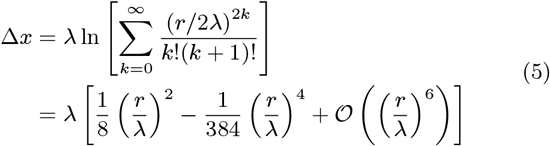

in absence of morphogen gradient variability and cell size variability (see Supplementary Material). For *r* = 2.45 μm and *λ* = 19.3 μm as found for SHH in the mouse neural tube [8], the shift is Δ*x* = 0.039 μm, or 0.8% of the cell diameter.

In the case of rectangular rather than circular cell areas, cells are confined to the interval [*x*_0_ – *r, x*_0_ + *r*]. The theoretically predicted shift is then approximately 0.052 μm in the mouse neural tube (see Supplementary Material) or 1% of the cell diameter. This agrees with the shift we measured in our simulations, Δ*x* = 0.0523 ± 0.0001 μm (mean ± SEM), confirming that spatial averaging of an exponential gradient results in a higher average concentration than centroid readout. Kinetic and area variability both increase Δ*x* (Fig. 3B), but it remains small enough (small fractions of a cell diameter) to be neglected in the analysis of tissue patterning under biological conditions where *r/λ* ≪ 1. Linear gradients [1] would not result in any shift at all.

### Spatial averaging barely reduces variability between gradients

Spatial and temporal averaging can reduce the positional error of morphogen gradients [32]. Previously, these mechanisms have been mainly analysed on the level of the morphogen readouts—typically transcription factors (TFs)—which are averaged by diffusion between nuclei [10, 33, 3, 16–19]. This is easily possible in a syncytium, as present in the early *Drosophila* embryo, but the role of TF diffusion in increasing patterning precision has remained controversial [34]. In an epithelium, nuclei are separated by cell membranes such that the averaging of morphogen-induced factors would require transport between cells, a complex and slow process with many additional sources of molecular noise [35, 36]. Epithelial cells potentially can, however, reap the benefits of spatial averaging by averaging the morphogen signal over their surface (Fig. 4A, green). Receptors may either be dispersed on the apical cell surface, or along the baso-lateral surface, or, in case of hormones, be limited to nuclei [37, 38]. In the latter case, morphogen receptors would be limited to a small patch, which could either be randomly positioned (Fig. 4A, blue), or located at the centroid of the cell (Fig. 4A, red). In the mouse neural tube, the SHH receptor PTCH1 is restricted to a cilium located on the apical surface [39]. The range of spatial averaging then depends on the cilium length and flexibility rather than the cross-sectional cell area (Fig. 4A, purple). We sought to analyse how the different spatial averaging strategies without cross-talk between neighbouring cells affect the variability of gradients, and thus the positional error.

Colours in panel 4A correspond to the colours in panels 4B– G). While the mean cell diameter *μ_δ_* greatly affects the gradient variability CV_*C*_, the readout strategy has only a moderate impact (Fig. 4B). The difference is most pronounced for large cells (*μ_δ_* = *μ_λ_*), where the sensed morphogen variability is largest if the cellular readout point is randomly placed (Fig. 4B, blue). Readout at the centroid or averaged over the entire cell yield similar sensed gradient variabilities. This can be understood since the theoretical considerations above predict only a small shift. Also, a cilium that averages the gradient concentration over larger regions than a single cell area barely reduces the sensed variability (Fig. 4C).

**Figure 4:**
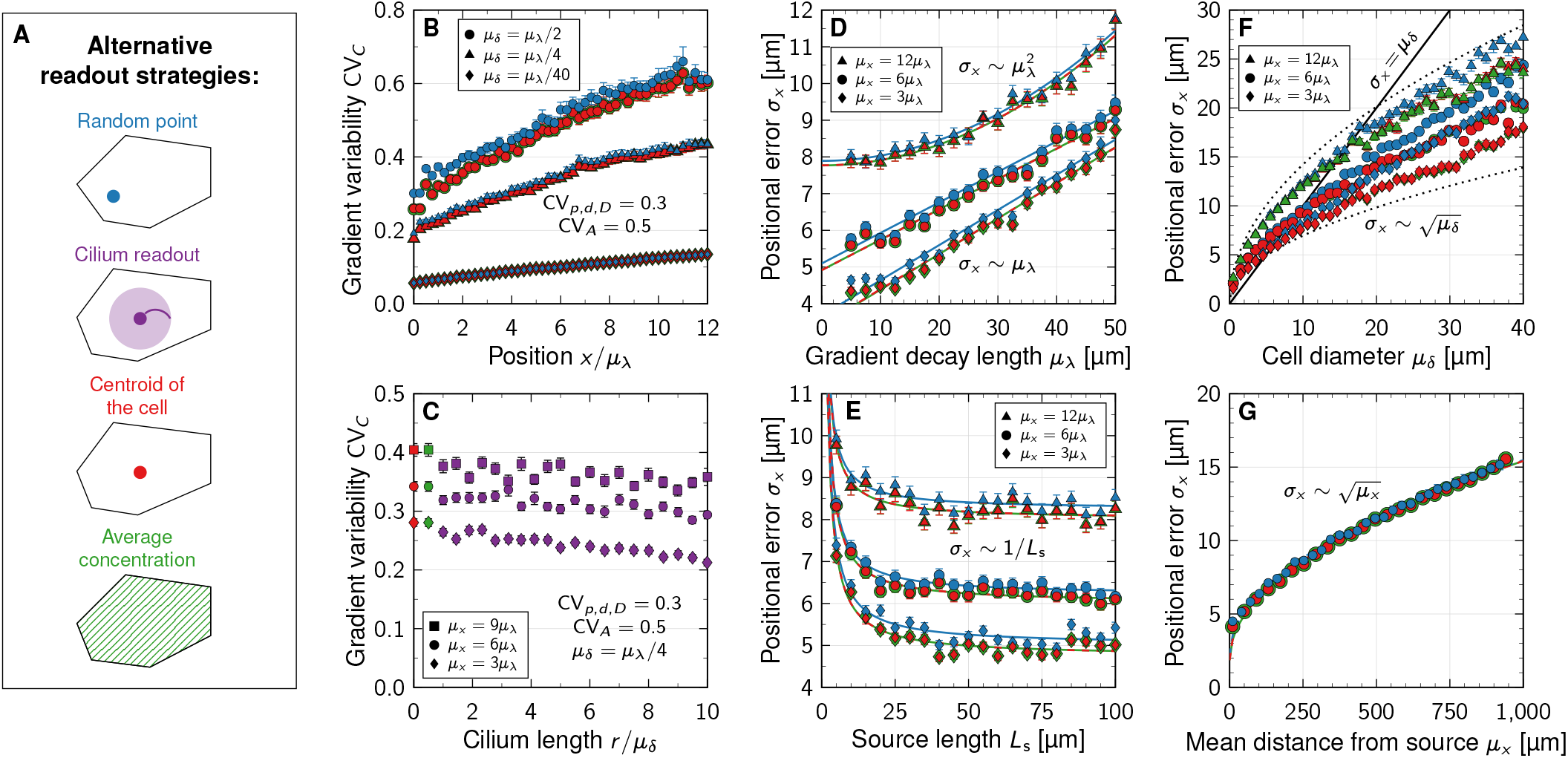
Impact of spatial averaging, gradient length, source size, cell diameter, and readout position on the positional error of morphogen gradients. **A** Four different methods how cells may read out morphogens. Colours in panels B–G correspond to these readout mechanisms. **B** The readout methods yield almost identical relative variability in the concentration over the patterning domain for small and medium sized cells. **C** Spatial averaging over a larger readout region (radius *r)* does not substantially decrease relative morphogen concentration variability. **D** Close to the source, the positional error scales linearly with the gradient decay length *μ_λ_*. Far in the domain, the scaling transitions to quadratic. **E** Asymptotically for short source length *L*_s_, the positional error is inversely proportional to 1/*L*_s_. **F** The positional error increases with the square root of the mean cell diameter *μ_δ_*. Dotted lines show the relationship 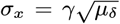 for *γ* = 2.2, 4.5 (lengths in units of μm). **G** Asymptotically, the positional error scales with the square root of the mean readout position *μ_x_*. Each data point in **B–G** corresponds to the mean ± SEM of *n* = 10^3^ independent simulations. Simulation parameters: *L*_p_ = 65*μ_δ_*, except in **G**; *μ_λ_* = 20 μm, except in **D**; *L*_s_ = 5*μ_δ_*, except in **E**; *μ_δ_* = 5 μm, CV_*p,d,D*_ = 0.3, CV_*A*_ = 0.5. See supplementary Table S1 for fit parameters.

In summary, larger cross-sectional cell diameters increase the variability of the morphogen concentration profiles, while spatial averaging over the cell surface barely reduces the gradient variability. Spatial averaging may, however, counteract detection noise at low morphogen concentrations far away from the source. It is currently unknown over which distance morphogen gradients operate. At distance 12*λ* from the source, for instance, exponential concentrations will have declined by *e*^12^ ≈ 160-thousand-fold. At such low levels, detection noise may dominate readout variability unless removed by spatial averaging.

### Scaling of the positional error with gradient length, source size, cell diameter and readout position

From dimensional analysis, the positional error of the gradient, *σ_x_*, being a measure of distance, must scale with a multiplicative combination of the length scales occurring in the patterning process. These can either originate from geometrical features of the tissue, or from the reaction-diffusion kinetics. We varied all relevant length scales in simulations and found that *σ_x_* is asymptotically proportional to the mean characteristic gradient decay length *μ_λ_* close to the source, but transitions to 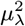 at larger distances (Fig. 4D). Additionally, it is inversely proportional to the source length *L*_s_, asymptotically for small *L*_s_ (Fig. 4E), but saturates for large sources. Moreover, the positional error increases with the square root of the mean cell diameter *μ_δ_* (Fig. 4F) and, up to an offset, with the square root of the mean position along the patterning axis *μ_x_* (Fig. 4G). Together, this can be expressed by the asymptotic scaling relationship

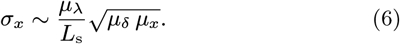

The linear dependency on the gradient length *μ_λ_* is due to the effect of gradient steepness on the positional error, and outweighs the reduction in gradient amplitude variability (Fig. 2H). It intuitively follows from *σ_x_* ≈ |*∂C*/*∂x*|^-1^ *σ_C_* ≈ *μ*_λ_CV_*C*_, which is a valid approximation when the average gradient has an exponential shape [24]. As before (Fig. 2F), at constant *μ_δ_*, a longer source reduces the gradient amplitude variability because noise is buffered by a larger number of source cells (Supplementary Material, Fig. S5). Narrower cells (smaller *μ_δ_*) reduce the positional error of the morphogen gradients according to the law of large numbers, 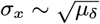. Cell width in the patterning domain is more influential than in the source, however, and the benefit of reducing cell width in the source alone is limited (Supplementary Material, Fig. S6). The deterministic limit (CV_*C*_ → 0, *σ_x_* → 0) is recovered in the continuum limit *μ_δ_* → 0. Domain boundaries can thus be defined more accurately at a certain target location *μx* within the tissue with narrow cells. Depending on the other lengths, the positional error can well be less than a cell diameter close enough to the source (Fig. 4F). We note that the previously reported linear scaling *σ_x_* ~ *μ_x_* [24] is valid only for idealized gradients that vary only through noise in *λ*, but not in their amplitude or from cell to cell. For the noisy, more physiological gradients simulated here, the positional error increases according to 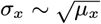 (asymptotically, Fig. 4G) and thus remains lower with increasing distance from the source than previously anticipated. This further challenges previous reports of excessive inaccuracy of the SHH and BMP gradients in the mouse neural tube [11].

### High precision of scaled patterns by parallel changes of gradient length, source size, and cell diameter in the *Drosophila* wing disc

The Decapentaplegic (Dpp) morphogen gradient in the *Drosophila* wing imaginal disc defines the position of several veins in the adult wing (Fig. 5A). Thus, the anterior-most limits of the Dpp source and the Dpp target gene spalt (*sal*) define the positions of the third (L3) and second (L2) longitudinal veins in the anterior compartment, respectively [40, 41], while the fifth longitudinal (L5) wing vein forms at the border between the expression domains of optomotor-blind (*omb*) and brinker (*brk*) in the posterior compartment [42]. The Dpp readout positions scale with the total length of the uniformly expanding patterning domain, such that the anterior position of the Sal-domain boundary remains roughly at 40–45% of the anterior domain length *L*_a_, while the posterior Omb domain boundary remains roughly at 50% of the posterior domain length *L*_p_ [41, 6, 43]. The gradient readout positions scale with the length of the patterning domain, because both the gradient length, *λ*, and the gradient amplitude *C*_0_ increase dynamically with the expanding tissue [6, 43–45] (Fig. 5B). On their own, the increases in *μ_λ_* and in *μ_x_* would lower the precision of the readout substantially over time (Eq. 6). However, the Dpp source widens in parallel, keeping the *μ*_λ_/*L*_s_ ratio at about 0.69 (Fig. 5B). Moreover, the apical cell diameter *μ_δ_* shrinks 3-fold close to the source from 4.5 to 1.5 μm [46–49, 27], which somewhat balances the increase in *μ_x_* over time. Plugging these dynamics into our model, the simulations showed that the positional error at *μ_x_* = 0.4*L*_a_ increases from 2.9 μm to 4.3 μm over developmental time (Fig. 5C, orange diamonds). If no compensation were taking place, the positional error would increase to about 6.5 μm in the same time period (Fig. 5C, blue circles).

The relative patterning precision, as quantified by the coefficient of variation CV_*x*_ = *σ_x_/μ_x_*, has even been reported to increase during development, as the CV of the distance between the L2 and L3 veins in the adult fly is only half (CV_*x*_ = 0.08) of that of the anterior-most Sal domain boundary (CV_*x*_ = 0.16) [41]. How this increase in precision is achieved has remained elusive. In light of Eq. 6, 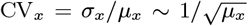 (Fig. 5D), such that the decreasing CV_*x*_ in adult stages could at least partly be a consequence of the increase in *μ_x_* = 0.4*L*_a_ between the stage when the precision of the Sal domain boundary was measured and the termination of Dpp-dependent patterning. The asymptotic relationship 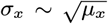 may thus provide an explanation of how the relative precision of patterning increases during *Drosophila* wing disc development.

### The effect of spatial correlation

Our theoretical considerations and simulations above are based on statistical independence between adjacent cells. To examine the effect of spatial correlations, we performed additional simulations in which this assumption was relaxed. We introduced a maximal degree of spatial correlation between neighbouring cells, given a certain degree of inter-cellular variability CV_*k*_, by sorting the kinetic parameters *p_i_*, *d_i_* and *D_i_* in ascending or descending order along the patterning axis after they had been drawn from their respective probability distributions, and then solved the reaction-diffusion problem (Eq. 2). The square-root increase of the positional error with the mean cell diameter remains intact in the presence of such spatial correlations between cells (see Supplementary Material, Fig. S3), with a slightly smaller prefactor. Since any physiological level of cell-to-cell correlation that preserves CV_*k*_ will lie somewhere between the uncorrelated and the maximally correlated extremes, the impact of such a form of spatial correlation on patterning precision can be expected to be minimal, and our findings remain valid also in presence of spatial correlations.

**Figure 5:**
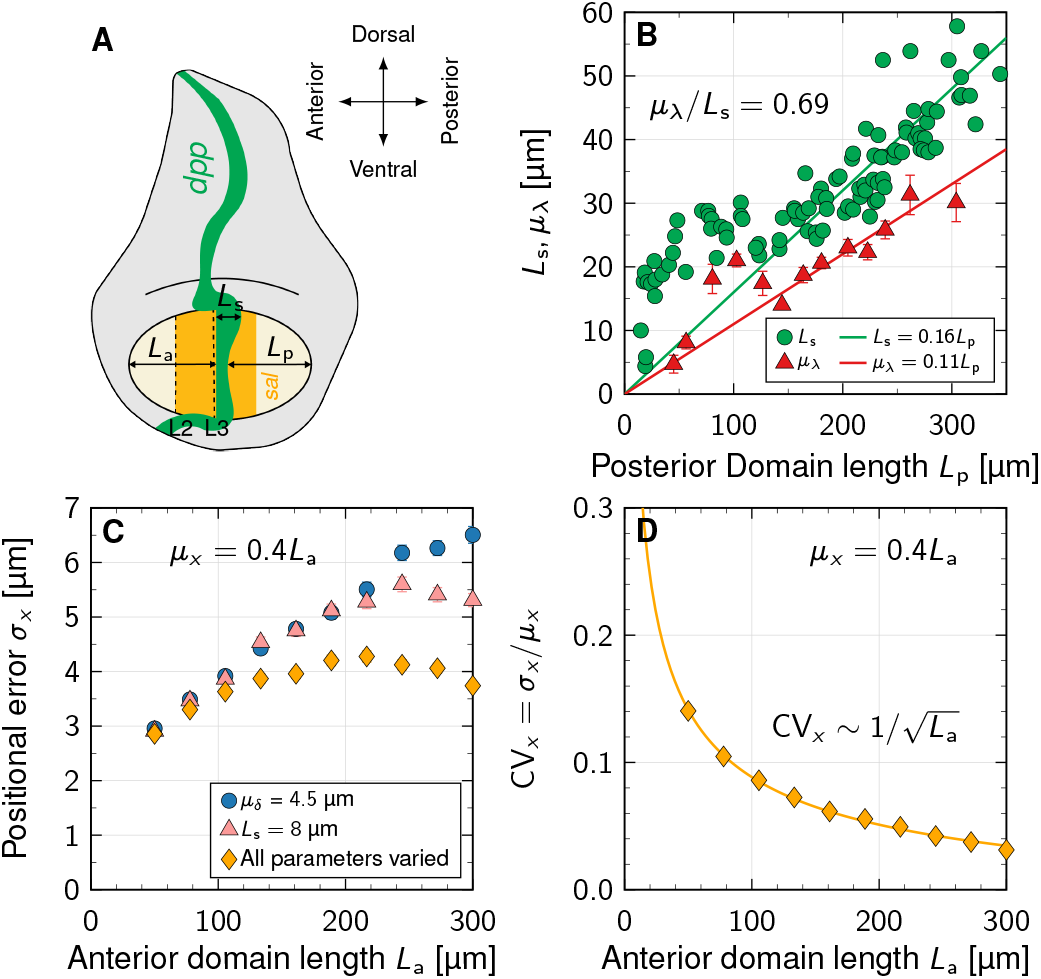
High precision of scaled patterns by parallel changes of gradient length, source size, and cell diameter in the *Drosophila* wing disc. **A** Schematic of Dpp-dependent patterning in the *Drosophila* wing disc. **B** The reported Dpp gradient length and source size increase in parallel with the expanding length, *L*_p_, of the posterior compartment. Data from Fig. S20 in [6]. **C** The predicted positional error at the relative readout position *μ_x_*/*L*_a_ = 40% is smallest when *μ_λ_* and *L*_s_ evolve according to the linear fits in B, and *μ_δ_* declines linearly from 4.5 to 1.5 μm (orange diamonds). For comparison, the positional error if *μ_δ_* is fixed and *μ_λ_*, *L*_s_ evolve (blue circles), or if the source length is fixed and *μ_λ_*, *μ_δ_* evolve (salmon triangles). **D** The predicted positional coefficient of variation CV_*x*_ = *σ_x_*/0.4*L*_a_ declines as the domain expands. See supplementary Table S1 for fit parameters.

An additional form of inter-cellular correlation may occur if nearby cells stem from the same lineage, and as such, may have correlated kinetic properties. In its most extreme form, neighbouring cells may share all their molecular parameters *p*, *d*, *D*, effectively becoming one wider joint cell in our model. We can use our results for cell autonomous noise to predict the dependency of patterning precision on the number of adjacent cells sharing their kinetic properties, *N*. Since the effective cell diameter simply becomes *N_μδ_*, the positional error will scale as 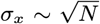. In this sense, the mean cell diameter *μδ* in our formulas may be interpreted as an effective spatial distance over which morphogen kinetics are shared, proportional to a spatial correlation length in the tissue, if any.

Cell-specific morphogen production and decay rates, and local variability in morphogen transport rates have not yet been quantified in epithelial tissues. A spatial coupling of molecular noise in dividing cells would require a perfectly symmetric division of cell contents upon cell division and the absence of cell-intrinsic noise. Dpp-containing endosomes are indeed distributed equally upon cell division in the *Drosophila* wing disc [50]. However, no cellular system without intrinsic noise has so far been reported. Differences between genetically identical sister cells were first shown for bacterial cells [51], but have since been demonstrated also for mammalian cells, and pose a key challenge in synthetic biology [52–54]. The coefficients of variation that we used are based on the reported variabilities of production and decay rates in single, genetically identical cells in cell culture [24].

There are further reasons why low spatial correlation of the kinetic parameters is to be expected. In pseudostratified epithelia, interkinetic nuclear migration (IKNM) introduces differences between cells as the cell cross-sectional areas change along the entire apical-basal axis over time [28]. As the tight junctions constitute a diffusion barrier between the apical and the baso-lateral domains, the apical receptor density between cells will change dynamically between cells if the apical receptor number is equal and fixed for all cells. To maintain the same receptor density, even though IKNM proceeds at different rates between neighboring cells, as reflected in the different nuclear positions along the apical-basal axis [28], the processes that balance receptor production and internalisation would need to be identical between neighboring cells, though differences in cell and nuclear volumes may also need to be compensated for. The same holds for the glyocalyx and extracellular matrix, which define the speed of morphogen diffusion, or fillipodia, in case of cytoneme-based transport. In summary, the combination of an unequal distribution of cell components in cell division, differences in the relative surface area to cell and nuclear volume, and intrinsic noise in gene expression must be expected to lead to individual differences between neighboring cells, even if they stem from the same lineage.

### Epithelial tissues patterned by morphogen gradients have small mean apical cell areas

After finding that patterning precision is greater with narrower cells in our model, we collected mean apical cell areas for a wide range of tissues from the literature to check whether cell diameters are small in tissues that rely on gradient-based patterning (Fig. 6). In the chick (cNT) and mouse neural tube (mNT) where SHH, BMP, and WNT gradients define the progenitor domain boundaries [55], the mean apical cell areas are largely around 7 μm^2^ and remain below 12 μm^2^ [48, 26, 29]. The chick embryonic ectoderm (cEE) appears to be patterned by BMP gradients [56], with mean apical cell area just below 12 μm^2^ [48]. In the *Drosophila* larval eye disc (dEYE), notum (dNP), and wing disc (dWL), Hedgehog (Hh), Decapentaplegic (Dpp), and Wg gradients pattern the epithelium [57, 58, 55], with mean apical cell areas smaller than 7 μm^2^ [47, 48, 46, 27]. The mean apical cell areas of the wing disc increase through the pre-pupal stages (dWP, dPW), to approximately 18 μm^2^ in the pupal stages [48, 27], other measurements in the *Drosophila* wing disc (dWD) report mean apical cell areas from 0 to 16 μm^2^ [46]. In the *Drosophila* eye antennal disc no gradient-based patterning was described (dEA folded; mean apical cell areas of approximately 33 μm^2^, dEA non-folded; with mean apical cell areas of approximately 39 μm^2^) [59]. For the peripodal membrane (dPE10–24) of the *Drosophila* eye disc, no gradientbased patterning has been described and mean apical cell areas range from 85 μm^2^ to more than 300 m^2^ [27]. In the *Drosophila* egg chamber (dEC), the mean apical cell areas decline from around 30 μm^2^ at stage 2/3 to around 10 μm^2^ by stage 6/7 [60], consistent with reported gradient-based patterning at stage 6 [61]; we did not find reports of earlier gradient-based patterning. While gradients pattern the *Drosophila* blastoderm syncytium [55], we are not aware of morphogen gradient readout during cellularisation. In the *Drosophila* embryo anterior pole (dEAP), the mean apical cell area is approximately 46 μm^2^ and in the embryo trunk (dET) roughly 35 μm^2^ [62], much larger than in the neural tube or wing disc. Before cellularisation, the situation is different from that in an epithelium in that free diffusion in the inter-nuclear space of the syncytium likely counteracts any sharp transition in the kinetic parameters as represented in our epithelial model, where cell membranes compartmentalise space. In the *Drosophila* L2 trachea (dL2T), no gradients have been reported and the mean apical cell areas are greater than 200 μm^2^ [63]. In the mouse embryonic lung (mLUNG), no morphogen gradients have been reported, despite chemical patterning [64]. The mean apical cell area is approximately 19 μm^2^ [65]. mean apical cell areas in the postnatal (P1–P21) cochlea are between 15 and 55 μm^2^ [66]. In adult mouse retinal pigment epithelial (mRPE) cells, the mean apical cell areas exceed 200 μm^2^ in young mice (P30) and increase to over 400 μm^2^ in old mice (P720) [67]. No gradient-based patterning was reported in mouse outer hair cells (mOHC1–3 P1,3,5,7.5); mean apical cell areas decrease from 35 μm^2^ (P1) to 16 μm^2^ (P7.5). No gradient-based patterning takes place in the inner hair cells (mIHC1 P1,3,6,7.5); mean apical cell areas decrease from 54 μm^2^ (P1) to 29 μm^2^ (P7.5) [66]. No gradient-based patterning was reported in the mouse ear epidermis (mEE), with mean apical cell areas of 1044 μm^2^ [68]. The data thus confirms that apical cell areas are small in tissues that employ gradient-based patterning. Our theory makes no prediction about the apical areas in tissues that do *not* employ gradient-based patterning, but in all cases that we have checked, apical areas are larger and appear to further increase in later developmental stages and in adult animals.

**Figure 6:**
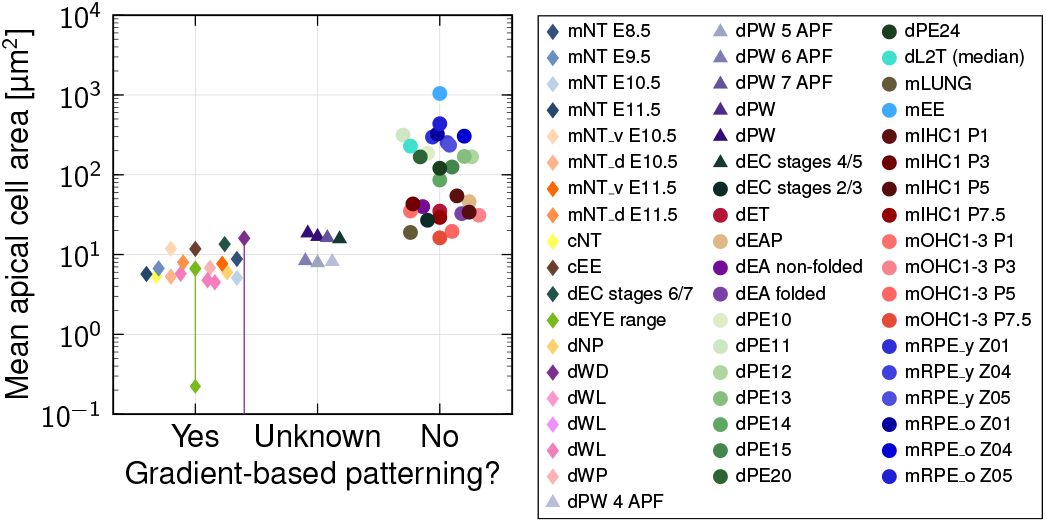
Epithelial tissues that use gradient-based patterning have small mean apical cell areas. For tissue abbreviations see main text.

## Discussion

We have shown that gradient precision decreases with increasing cross-sectional area of the patterned cells. Consistent with our prediction, apical surface areas are small in epithelia that employ gradient-based patterning. In curved domains, spatial precision will be higher on the inside, where the average cell diameter is smaller. In the mouse neural tube, the SHH-sensing cilium is indeed located on the inner, apical surface [39], while in the flat *Drosophila* imaginal discs, cells sense Hedgehog along the entire apical-basal axis [69]. In the *Drosophila* wing disc, the apical cell diameters shrink in the center of the domain, such that the apical areas are almost twofold smaller close to the source, and increase roughly linearly [47, 70, 49, 71]. In the eye disc, the size gradient is even more pronounced, with tiny apical areas in the Dpp secreting morphogenetic furrow [47]. The declining apical cell diameters have previously been accounted to a mechanical pressure feedback caused by growth [72, 46]. However, signalling by Dpp, the fly homolog of mammalian BMP2/4, has been shown to result in taller cells with smaller cross-sectional area in its patterning domain compared to other parts of the *Drosophila* wing and eye disc [47, 70, 49, 71]. Similarly, the morphogens SHH and WNT have been observed to increase cell height and reduce the cell cross-sectional area via their impact on actin polymerisation, myosin localisation and activity in the embryonic mouse neural tube and lung [65, 70, 73–75]. In light of our study, it is possible that the morphogen-dependent reduction in the cross-sectional cell area via positive modulation of cell height serves to enhance patterning precision. The precision advantage of small cell diameters may then have led to the emergence of pseudostratification in epithelial monolayers, a phenomenon that has so far remained unexplained. Our finding that wide cells and very large cell area variability are both detrimental to patterning precision indicate that there is potentially a window for epithelial pseudostratification in which patterning precision is optimal: High cell density benefits precision because cell diameters are small, but with nuclei much wider than the average cell diameter [28], precision would decline due to large area variability. It is remarkable that all tissues that we analysed seem to lie in the optimal range of this trade-off [27]. This aspect deserves further research and needs to be tested with additional experiments.

We have revealed scaling relationships between the positional error, cell diameter, gradient decay length and source length (Eq. 6). In follow-up work, we found that they also hold for non-exponential gradients arising from non-linear morphogen degradation [76], as far as they were studied. These relationships predict that morphogen gradients remain highly accurate over very long distances, providing precise positional information even far away from the morphogen source. Our results are system-agnostic, and could thus apply widely in development. The compensation between cell diameter, gradient length, source size and readout location, which we have found here, allows a patterning system to tune its length scales to achieve a particular level of spatial precision. Our theoretical work suggests a potential evolutionary benefit for a developmental mechanism that regulates features such as the cell diameter or the *λ/L*s ratio to maintain high patterning precision. A loss in precision due to a shift in readout position away from the morphogen source, for instance, can be compensated for by narrower cells in the source or in the patterning domain. This allows developmental systems to maintain high patterning precision at readout positions that scale with a growing tissue domain.

Whether pre-steady-state gradients, as likely play a role in the patterning of the *Drosophila* wing disc [44], follow the same behavior as discovered here for the steady state, remains an open question for future research. Assuming that they do, our results offer a potential explanation for the observed increase in relative patterning precision during wing disc development.

## Methods

### Generation of variable morphogen gradients

The patterning axis was constructed as follows: A random cell area *A_i_* was drawn for cell *i* = 1, and then converted to a diameter 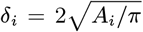, which assumes that cell surfaces are roughly isotropic. This process was repeated for the next cells *i* = 2, 3,… until their cumulated diameters matched the domain length *L*_s_ or *L*_p_. To control the mean cell diameter *μ_δ_*, cell areas were drawn with a mean value of 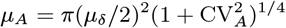 for given *μ_δ_* and CV_*A*_, as follows from the transformation properties of log-normal random variables, such that indeed 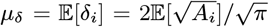. The patterning axis was then discretized into subintervals of length *δ_i_*, the source and patterning domains were pasted together such that *x* = 0 marked the source boundary, and random kinetic parameters *p_i_, d_i_, D_i_* were drawn independently for each cell from log-normal distributions. Note that the results reported in this work are largely independent of the specific choice of probability distribution, given that they do not allow for very small (or even negative) kinetic parameters, which would not be compatible with a successful morphogen transport and patterning process. A gamma distribution with the same mean and variance, for example, yields largely unchanged behavior (see Supplementary Material, Fig. S4).

We then solved Eq. 2 numerically on the discretized domain using Matlab’s built-in fourth-order boundary value problem solver `bvp4c` (version R2020b). Continuity of the morphogen concentration and its flux was imposed at each cell boundary. Further technical details can be found in [24]. Each simulation was repeated *n* = 10^3^ times with independent random parameters and cell areas.

### Gradient parameter extraction

We determined the amplitude *C*_0_ and decay length *λ* for each numerically generated noisy morphogen gradient by fitting the deterministic solution to it. With no-flux boundaries, the gradient shapes are hyperbolic cosines that slightly deviate from a pure exponential in the far end [24]. We fitted these inside the patterning domain to obtain *C*_0_ and *λ* after logarithmisation of the morphogen concentration as detailed in [24].

Since the fitted characteristic gradient length *λ* drifts away from the prescribed value for noisy gradients depending on which of the kinetic parameters is varied and by how much

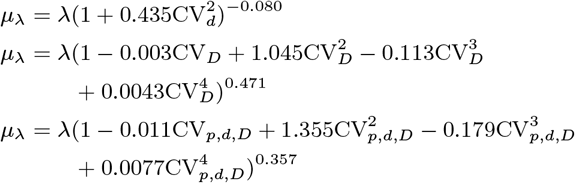

[24], we corrected for this drift in our numerical implementation to be able to use the true observed value of *μ_λ_* in our results: where *λ* is the deterministic (prescribed) value. When only the production rate *p* was varied, *μ_λ_* = *λ*. These empirical relationships approximate the data shown in Fig. 8G in [24].

## Code Availability

The source code is released under the 3-clause BSD license. It is available as a public git repository at https://git.bsse.ethz.ch/iber/Publications/2022_adelmann_vetter_cell_size.

## Acknowledgements

We thank Marco Meer for providing cell area data, and Fernando Casares and Nikolaos Doumpas for discussions. This work was funded by SNF Sinergia grant CRSII5_170930.

## Author Contributions

RV & DI conceived the study, JA and RV developed the numerical framework and produced the figures, JA carried out the simulations and analysis, RV developed the theory, DI contributed the supporting data. All authors wrote the manuscript.

