## Supplemental Material for "Impact of cell size on morphogen gradient precision"

### Impact of cell size on morphogen gradient precision: Supplementary material

In this supplementary document, we theoretically show that averaging morphogen concentrations over a spatial region (such as cell areas) can shift the effective readout position compared to point-like readout, and we derive the corresponding shift $\Delta x$ analytically for isotropic and for rectangular cell shapes. We focus on exponential morphogen gradients here as they arise in systems with diffusion-driven morphogen transport and uniform linear degradation, but note that the developed formalism can be applied directly also to other gradient shapes. Moreover, the impact of spatial correlation of the kinetic cell parameters on the positional error, the choice of the kinetic parameter distribution and the effect of cell number in the source domain are discussed.

#### Readout in a continuous domain

Consider an exponential morphogen concentration gradient

$$C(x) = C_0 \exp\left[-\frac{x}{\lambda}\right]$$

with concentration $C_0$ at the source at $x = 0$, and decay length $\lambda$. Assuming a continuous readout based on a threshold concentration $C_\theta = C(x_\theta)$, a positional identity boundary forms at position

$$x_\theta = \lambda \ln\left[\frac{C_0}{C_\theta}\right]. \tag{S1}$$

This mechanism allows for gradient-based tissue patterning, where individual patterning domains are delineated by different boundary positions $x_\theta$ resulting from different readout thresholds $C_\theta$.

#### Readout in a tissue of isotropic cells

For morphogen readout in a cellular tissue, we consider several different cases in a unified description. Cells can either sense the morphogen concentration at a singular point, averaged over a spatial region with radius $r$ about that point (which may or may not be smaller than a cell), or as an average concentration over the entire cell area. We denote this readout region by $\Omega$ (Fig. S1). The average concentration in $\Omega$ is

$$\langle C\rangle = \frac{\int_\Omega C(x)\, d\Omega}{\int_\Omega d\Omega}.$$

Assuming that the averaging domain is circular (i.e., the cell areas have no orientational bias) in a two-dimensional tissue cross section or surface, we can approximate $\Omega$ as a disk with radius $r$ about a center point $(x_0, 0)$:

$$\Omega = \left\{(x,y) \mid (x-x_0)^2 + y^2 < r^2\right\}.$$

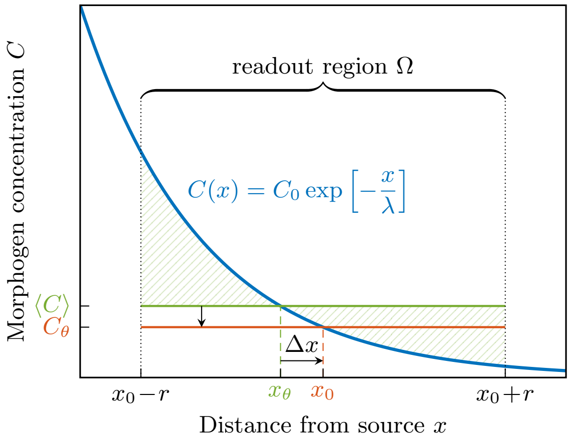


**Figure S1:** Averaging an exponential morphogen concentration (blue) over a local region such as the cell area leads to a larger readout concentration (green) than taking the concentration at the middle of the region (red). To compensate for this effect, the readout position shifts downhill (away from the source) by a distance $\Delta x$ from $x_\theta$ to $x_0$.

In the case where the concentration is averaged over the entire cell area, $r$ is the effective cell radius. The average concentration thus becomes

$$\begin{aligned}\langle C\rangle &= \frac{C_0}{\pi r^2}\int_\Omega \exp\left[-\frac{x}{\lambda}\right] d\Omega \\ &= \frac{C_0}{\pi r^2} 2\pi r\lambda \exp\left[-\frac{x_0}{\lambda}\right] I_1\left(\frac{r}{\lambda}\right)\end{aligned}$$

where

$$I_n(z) = \sum_{k=0}^{\infty} \frac{(z/2)^{2k+n}}{k!(k+n)!}$$

is the modified Bessel function of the first kind for integer $n$. The series converges very quickly if $r \ll \lambda$, such that higher order terms in $r/\lambda$ can be dropped. Substitution and expansion of the Bessel function yields

$$\begin{aligned}\langle C\rangle &= C(x_0)\frac{2\lambda}{r} I_1\left(\frac{r}{\lambda}\right) \\ &= C(x_0)\sum_{k=0}^{\infty}\frac{(r/2\lambda)^{2k}}{k!(k+1)!} \\ &= C(x_0)\left[1 + \frac{1}{8}\left(\frac{r}{\lambda}\right)^2 + \frac{1}{192}\left(\frac{r}{\lambda}\right)^4 + \mathcal{O}\left(\left(\frac{r}{\lambda}\right)^6\right)\right].\end{aligned}$$

Thus, the mean concentration $\langle C\rangle$ is larger than the one in the middle of the readout domain, $C(x_0)$, and this deviation increases with larger readout regions and shorter gradient decay lengths.

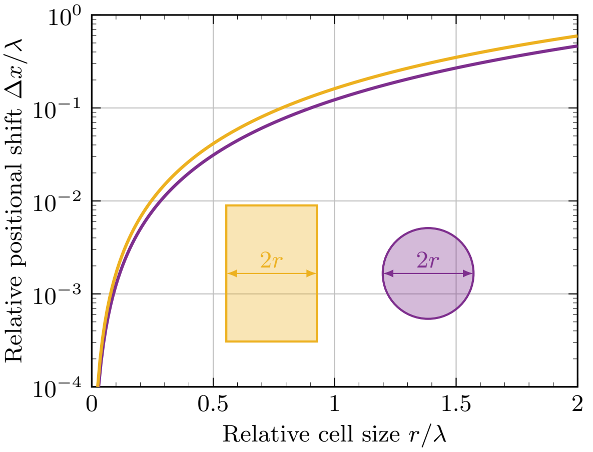


**Figure S2:** Readout boundary shift due to spatial averaging as a function of the size over which the morphogen concentration is averaged. The purple line shows the isotropic case with a circular averaging region (Eq. S2); the orange line represents the case with rectangular cells (Eq. S4).

If threshold-based readout operates on the averaged concentration, we must have $C_\theta = \langle C \rangle$. Therefore,

$$\frac{C(x_\theta)}{C(x_0)} = \exp\left[-\frac{x_\theta - x_0}{\lambda}\right] = \sum_{k=0}^{\infty} \frac{(r/2\lambda)^{2k}}{k!(k+1)!}.$$

The location of domain boundaries is shifted down the concentration gradient by the distance

$$\Delta x = x_0 - x_\theta = \lambda \ln\left[\sum_{k=0}^{\infty} \frac{(r/2\lambda)^{2k}}{k!(k+1)!}\right] \tag{S2}$$

as shown in Fig. S1. Notably, the shift is independent of both the gradient amplitude $C_0$ and the concentration threshold $C_\theta$ for an exponential gradient. Therefore, it is the same for all readout positions in the pattern if the averaging radius $r$ and the decay length $\lambda$ are spatially invariant, such that all domain boundaries are shifted equally by this averaging effect. Eq. S2 is plotted in Fig. S2.

Using the power series expansion of the natural logarithm,

$$\ln[1+x] = \sum_{k=1}^{\infty} (-1)^{k+1} \frac{x^k}{k} = x - \frac{x^2}{2} + \mathcal{O}\left(x^3\right),$$

the boundary shift can be expanded to

$$\Delta x = \lambda \left[\frac{1}{8}\left(\frac{r}{\lambda}\right)^2 - \frac{1}{384}\left(\frac{r}{\lambda}\right)^4 + \mathcal{O}\left(\left(\frac{r}{\lambda}\right)^6\right)\right].$$

For a mean cell radius of $r = 2.5\,\mu\text{m}$ and a gradient decay length of $\lambda = 20\,\mu\text{m}$, the shift is $\Delta x \approx 0.039\,\mu\text{m}$.

By combining Eqs. S1 and S2, we find the mean domain boundary position at

$$x_0 = x_\theta + \Delta x = \lambda \ln\left[\frac{C_0}{C_\theta} \sum_{k=0}^{\infty} \frac{(r/2\lambda)^{2k}}{k!(k+1)!}\right]. \tag{S3}$$

#### Readout in a tissue of rectangular cells

We now derive the downhill shift $\Delta x$ also for rectangular cell areas, effectively rendering the problem one-dimensional. This scenario corresponds to a tissue composed of cuboidal cells in which the morphogen gradient forms in a direction perpendicular to one of the cells' axes. In this case,

$$\Omega = \{(x,y) \mid |x - x_0| < r\}.$$

Averaging over the cell area thus gives

$$\langle C \rangle = \frac{C_0}{2r} \int_\Omega \exp\left[-\frac{x}{\lambda}\right] d\Omega = C(x_0) \frac{\lambda}{r} \sinh\left[\frac{r}{\lambda}\right]$$

Requiring again that the readout threshold be the average concentration, $C_\theta = \langle C \rangle$, yields

$$\frac{C(x_\theta)}{C(x_0)} = \exp\left[-\frac{x_\theta - x_0}{\lambda}\right] = \frac{\lambda}{r} \sinh\left[\frac{r}{\lambda}\right].$$

The shift in the readout position then follows as

$$\Delta x = x_0 - x_\theta = \lambda \ln\left[\frac{\lambda}{r} \sinh\left(\frac{r}{\lambda}\right)\right] \tag{S4}$$

which expands to

$$\Delta x = \lambda \left[\frac{1}{6}\left(\frac{r}{\lambda}\right)^2 - \frac{1}{180}\left(\frac{r}{\lambda}\right)^4 + \mathcal{O}\left(\left(\frac{r}{\lambda}\right)^6\right)\right].$$

Eq. S4 is plotted in Fig. S2. For a mean cell radius of $r = 2.5\,\mu\text{m}$ (which in this case corresponds to the half-width of the rectangular cells) and a gradient decay length of $\lambda = 20\,\mu\text{m}$, the shift is $\Delta x \approx 0.052\,\mu\text{m}$.

In analogy to Eq. S3, the mean domain boundary position is found at

$$x_0 = x_\theta + \Delta x = \lambda \ln\left[\frac{C_0}{C_\theta} \frac{\lambda}{r} \sinh\left(\frac{r}{\lambda}\right)\right].$$

in tissues composed of rectangular cells.

#### Impact of spatial correlation on the positional error

In the main article, we assumed uncorrelated morphogen kinetics. Here, we demonstrate how spatial correlation affects the positional error. First, we consider total correlation, where all three kinetic parameters ($p$, $d$, $D$) are the same for all cells, but are still varied between different simulations (different tissues). In this limiting case, morphogen gradient variability occurs only between tissues, not within them. The positional error is significantly greater than with independent cells, and the square-root scaling is lost (Fig. S3, green triangles), because to the morphogen gradient, the tissue effectively appears like a homogeneous continuum with uniform properties.

Next, we consider, as a second extreme case, a maximal degree of cell-to-cell correlation in the kinetic parameters, while preserving their probability distributions within the tissue. The kinetic cell parameters ($p_i$, $d_i$, $D_i$) are drawn individually and independently for each cell, but are then sorted along the patterning axis and assigned to the cells $i$, prior to solving the reaction-diffusion equation. Sorting does not affect the patterning precision appreciably, independent of the ordering (Fig. S3). In comparison to zero correlation, sorting slightly reduces the positional error—an effect that is most pronounced for larger cell diameters. But even with this maximal level of spatial cell-to-cell correlation, the square-root scaling of the positional error holds. Intermediate levels of spatial correlation can be expected to yield positional errors lying in between the curves for zero and maximal cell-to-cell correlation.

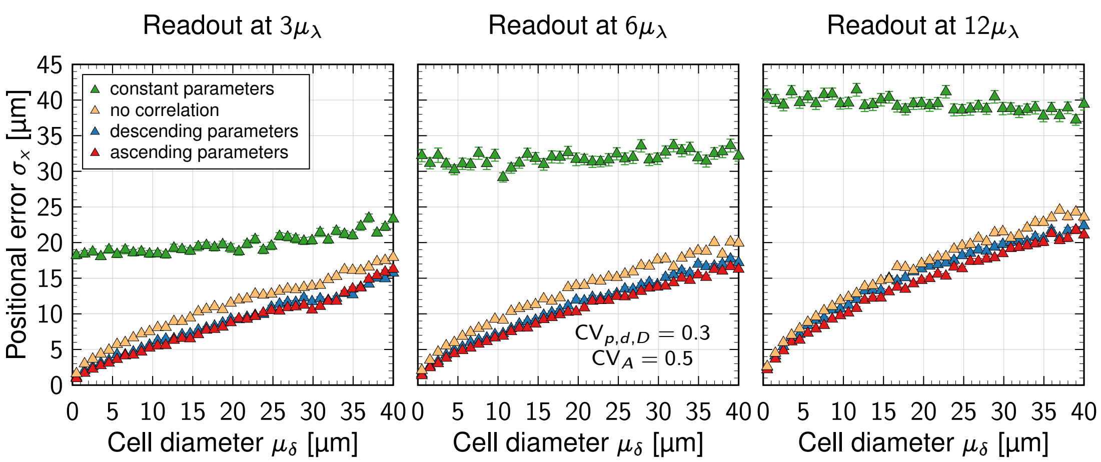


**Figure S3: Impact of correlation on the positional error at different readout positions and cell diameters.** Green triangles represent total correlation (all cells have equal kinetic parameters), yellow triangles represent no correlation (as presented in the main Fig. 4F). Blue (red) triangles correspond to the case with maximal spatial correlation at given cell-to-cell variability $\mathrm{CV}_{p,d,D}$, where the cell parameters were drawn from log-normal distributions and then sorted in descending (ascending) order. All simulations were repeated $n = 10^3$ times and the mean positional error ± SEM is plotted.

#### Choice of the kinetic parameter distribution

In the main article, we assumed log-normally distributed morphogen kinetics. In this section, we show that our results are largely independent of the probability distribution assumed for the kinetic parameters, provided that it meets certain physiological criteria:

- The morphogen production rates, degradation rates and diffusivities must be strictly positive. This rules out a normal distribution.
- The probability density of near-zero kinetic parameters must vanish quickly, as otherwise no successful patterning can occur. For example, a tiny diffusion coefficient would not enable morphogen transport over biologically useful distances within useful time periods. This rules out a normal distribution truncated at zero, because very low diffusivities would occur rather frequently for such a distribution.

We repeated the simulations shown in Figs. 2A,B and 4F with a gamma distribution in place of the log-normal distribution. Among other distributions that are conceivable, a gamma distribution with appropriate shape parameter $\alpha$ and inverse scale parameter $\beta$ fulfills the above criteria. In order to recover the mean and variance of the kinetic parameters, we set $\alpha_k = 1/\mathrm{CV}_k^2$ and $\beta_k = \mathrm{CV}_k^2/\mu_k$, where $\mathrm{CV}_k$ is the coefficient of variation and $\mu_k$ the mean value of a specific kinetic parameter $k$. As can be appreciated from Fig. S4, the results are not significantly altered by the specific choice of probability distribution, and our conclusions remain valid. The scaling exponents are consistent within statistical errors.

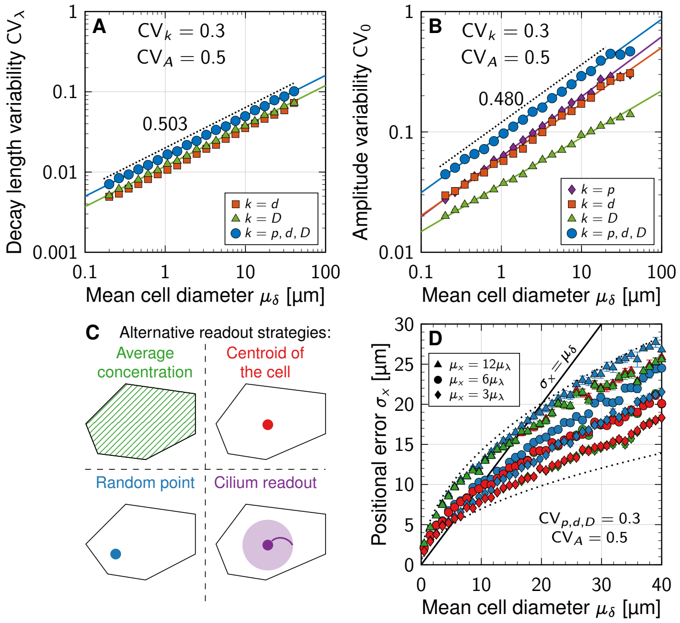


**Figure S4: Gradient variability and positional error under gamma-distributed morphogen kinetics.** All simulations were repeated $n = 10^3$ times and the mean values ± SEM are plotted. **A,B** The same scaling laws for the gradient variability found for the gamma and log-normal distributions (Fig. 2A,B) are consistent. **C** Different readout strategies (identical to Fig. 4A). **D** Square-root scaling of the positional error with the cell diameter is found also with gamma-distributed morphogen kinetics. Symbol colours in D correspond to the different morphogen sensing strategies in C.

#### Effect of cell number in the source domain on gradient precision

In the main article, we showed that patterning precision increases with narrower cells and wider sources. These effects are coupled—wider sources will be composed of more cells if the average cell diameter remains constant. In this section, we demonstrate that the positional error is mainly dominated by the cell diameter rather than the source size, and that the found scaling $\sigma_x \sim 1/L_\mathrm{s}$ (Eq. 6) is largely due to higher cell numbers in wider sources.

Increasing the number of cells in a source of fixed length improves the precision of the morphogen gradient parameters according to the asymptotic relationship

$$\mathrm{CV}_{\lambda,0} \sim \sqrt{\frac{\mu_{\delta_\mathrm{s}}}{L_\mathrm{s}}} \sim \sqrt{\frac{1}{N_\mathrm{cells}}},$$

where $N_\mathrm{cells}$ is the number of cells in the source domain (Fig. S5A,B). They thus approximately follow the law of large numbers. The positional error decreases analogously with increased cell number in a source of fixed length (Fig. S5C). If, on the other hand, the number of source cells is fixed but the

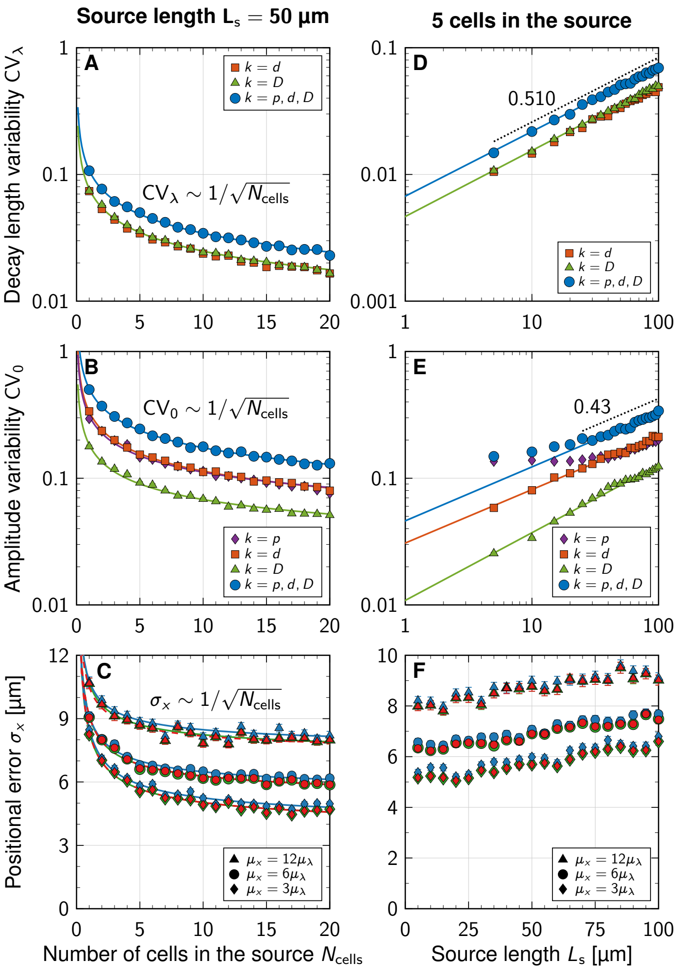


**Figure S5: Effect of source length and number of source cells on gradients precision. A,B** In a source of fixed length $L_s$, there is less variability in the gradient parameters $\lambda$ and $C_0$ as the number of constituting cells increases. **C** The positional error decreases with more cells in a source of fixed length, but saturates beyond about 5 source cells. **D,E** The gradient parameters become more variable in wider sources consisting of a fixed number of cells. **F** The positional error mildly increases in wider morphogen sources with fixed cell count. Colours in C,F correspond to readout strategies shown in Fig. S4C. All data points show mean values ± SEM from $n = 10^3$ simulations. Model parameters: $\mu_{\delta_p} = 5$ μm, $CV_{p,d,D} = 0.3$, $CV_A = 0.5$, $\mu_\lambda = 20$ μm.

source size increases, the variability in the gradient parameters increases according to power laws (Fig. S5D,E),

$$CV_{\lambda,0} \sim \mu_\delta^\alpha \quad \text{and} \quad CV_0 \sim \mu_\delta^\beta \tag{S5}$$

with exponents $\alpha = 0.510 \pm 0.005$ (Fig. S5D, blue curve) and $\beta = 0.43 \pm 0.02$ (Fig. S5E, blue curve), suggesting again $CV_{\lambda,0} \sim \sqrt{\mu_{\delta_s}/L_s}$. A source composed of a fixed number of cells yields only a mildly greater positional error if its constituent cells have a larger average diameter, however (Fig. S5F). In these simulations, the mean cell diameter in the patterning domain was fixed. Thus, in order to achieve high spatial gradient precision, a morphogen source must have a large number of cells with small diameters, but the cell count is more decisive than the source length.

To study the competition of cell sizes between the source and patterning domain, we then changed the mean cell diameter separately in both subdomains, retaining the mean diameter in the other at a constant value. No further appreciable increase in gradient precision takes place once the mean cell diameter in the source subceeds the one in the patterning domain ($\mu_{\delta_s} < \mu_{\delta_p}$, Fig. S6). The mean cell diameter in the source has a limited impact on gradient precision (Fig. S6, pink symbols) compared to the mean diameter in the patterning domain (Fig. S6, yellow symbols). Overall, this suggests that a large number of narrow cells in both the source and patterning domain, but mainly in the latter, is advantageous for patterning precision.

#### Fit parameters

In Table S1, we list all functional relationships used to fit the data shown in the main article and this supplementary document, together with the fit parameters and their standard errors (SE).

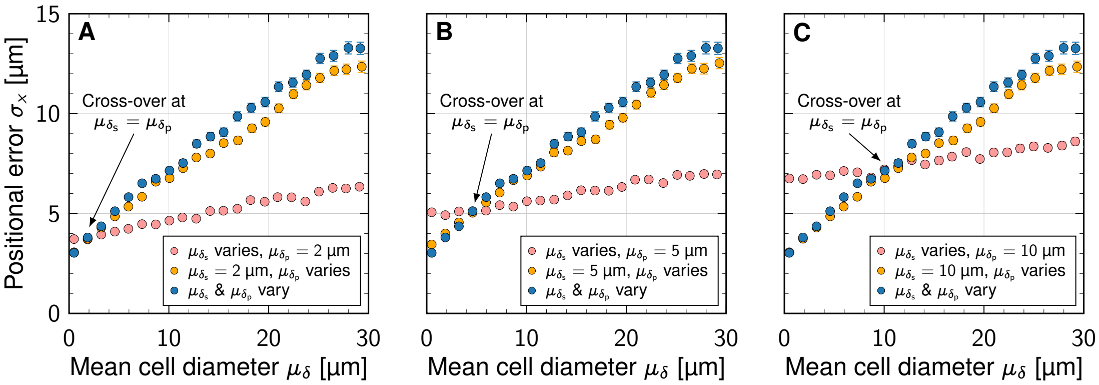


**Figure S6: Separate effects of the mean cell diameter in the source and patterning domains on the positional error. A,B,C** Change of positional error at $\mu_x = 3\mu_\lambda = 60$ μm, as the mean cell diameter is varied only in the source ($\mu_\delta = \mu_{\delta_s}$, pink), only in the pattern ($\mu_\delta = \mu_{\delta_p}$, yellow) or in both simultaneously ($\mu_\delta = \mu_{\delta_s} = \mu_{\delta_p}$, blue), but is fixed elsewhere (at 2, 5, 10 μm in A, B, C, respectively). All simulations were repeated $n = 10^3$ times and the mean values ± SEM are plotted. Model parameters: $L_s = 5\mu_{\delta_s}$, $CV_{p,d,D} = 0.3$, $CV_A = 0.5$.

**Table S1:** Summary of the fit functions and their parameters. All lengths are in micrometres.

| Figure | Model | Legend entry | a | SE(a) | b | SE(b) |
|---|---|---|---|---|---|---|
| 2A | $\ln \mathrm{CV}_\lambda = a \ln \mu_\delta + b$ | $k = D$ | 0.507 | 0.002 | −4.528 | 0.004 |
| | | $k = p, d, D$ | 0.510 | 0.004 | −4.199 | 0.008 |
| 2B | $\ln \mathrm{CV}_0 = a \ln \mu_\delta + b$ | $k = p$ | 0.497 | 0.003 | −4.528 | 0.004 |
| | | $k = d$ | 0.457 | 0.004 | −2.847 | 0.006 |
| | | $k = D$ | 0.387 | 0.006 | −3.403 | 0.008 |
| | | $k = p, d, D$ | 0.472 | 0.005 | −2.396 | 0.007 |
| 2E | $\mathrm{CV}_\lambda = b$ | $k = D$ | — | — | 0.0249 | 0.0001 |
| | | $k = p, d, D$ | — | — | 0.0343 | 0.0001 |
| 2F | $\mathrm{CV}_0 = a/L_\mathrm{s} + b$ | $k = p$ | 1.087 | 0.038 | 0.095 | 0.002 |
| | | $k = D$ | −0.158 | 0.010 | 0.070 | 0.001 |
| | | $k = p, d, D$ | 0.870 | 0.025 | 0.160 | 0.001 |
| 2G | $\mathrm{CV}_\lambda = b$ | $k = d$ | — | — | 0.0238 | 0.0002 |
| | | $k = D$ | — | — | 0.0246 | 0.0001 |
| | | $k = p, d, D$ | — | — | 0.0338 | 0.0001 |
| 4D | $\sigma_x = a\mu_\lambda + b$ | $\mu_x = 3\mu_\lambda$ average | 0.097 | 0.004 | 3.4 | 0.1 |
| | | $\mu_x = 3\mu_\lambda$ centroid | 0.087 | 0.004 | 3.4 | 0.1 |
| | | $\mu_x = 3\mu_\lambda$ random | 0.096 | 0.004 | 3.7 | 0.1 |
| | | $\mu_x = 6\mu_\lambda$ average | 0.083 | 0.003 | 4.9 | 0.1 |
| | | $\mu_x = 6\mu_\lambda$ centroid | 0.083 | 0.003 | 4.9 | 0.1 |
| | | $\mu_x = 6\mu_\lambda$ random | 0.083 | 0.003 | 5.1 | 0.1 |
| 4D | $\sigma_x = a\mu_\lambda^2 + b$ | $\mu_x = 12\mu_\lambda$ average | 0.0014 | 0.0001 | 7.8 | 0.1 |
| | | $\mu_x = 12\mu_\lambda$ centroid | 0.0014 | 0.0001 | 7.8 | 0.1 |
| | | $\mu_x = 12\mu_\lambda$ random | 0.0014 | 0.0001 | 7.9 | 0.1 |
| 4E | $\sigma_x = a/L_\mathrm{s} + b$ | $\mu_x = 3\mu_\lambda$ average | 12.5 | 0.9 | 4.75 | 0.05 |
| | | $\mu_x = 3\mu_\lambda$ centroid | 12.6 | 0.8 | 4.74 | 0.05 |
| | | $\mu_x = 3\mu_\lambda$ random | 12.3 | 1.0 | 5.01 | 0.05 |
| | | $\mu_x = 6\mu_\lambda$ average | 11.4 | 0.6 | 6.01 | 0.03 |
| | | $\mu_x = 6\mu_\lambda$ centroid | 11.3 | 0.6 | 6.01 | 0.03 |
| | | $\mu_x = 6\mu_\lambda$ random | 10.9 | 0.6 | 6.20 | 0.03 |
| | | $\mu_x = 12\mu_\lambda$ average | 8.9 | 1.0 | 8.01 | 0.06 |
| | | $\mu_x = 12\mu_\lambda$ centroid | 8.9 | 1.0 | 8.01 | 0.06 |
| | | $\mu_x = 12\mu_\lambda$ random | 8.5 | 1.0 | 8.24 | 0.05 |
| 4G | $\sigma_x = a\sqrt{\mu_x} + b$ | average | 0.429 | 0.003 | 1.86 | 0.06 |
| | | centroid | 0.429 | 0.003 | 1.85 | 0.06 |
| | | random | 0.421 | 0.003 | 2.17 | 0.07 |
| 5D | $\mathrm{CV}_x = a/\sqrt{L_\mathrm{p}} + b$ | | 1.28 | 0.02 | −0.039 | 0.002 |
| S5A | $\mathrm{CV}_\lambda = a/\sqrt{N_\mathrm{cells}}$ | $k = D$ | 0.0778 | 0.0006 | — | — |
| | | $k = p, d, D$ | 0.1082 | 0.0005 | — | — |
| S5B | $\mathrm{CV}_0 = a/\sqrt{N_\mathrm{cells}} + b$ | $k = p$ | 0.293 | 0.009 | 0.019 | 0.004 |
| | | $k = d$ | 0.325 | 0.003 | 0.011 | 0.001 |
| | | $k = D$ | 0.171 | 0.004 | 0.014 | 0.002 |
| | | $k = p, d, D$ | 0.490 | 0.006 | 0.019 | 0.003 |
| S5C | $\sigma_x = a/N_\mathrm{cells} + b$ | $\mu_x = 3\mu_\lambda$ average | 4.9 | 0.2 | 3.48 | 0.06 |
| | | $\mu_x = 3\mu_\lambda$ centroid | 4.9 | 0.2 | 3.47 | 0.06 |
| | | $\mu_x = 3\mu_\lambda$ random | 4.8 | 0.2 | 3.73 | 0.07 |
| | | $\mu_x = 6\mu_\lambda$ average | 4.2 | 0.1 | 4.94 | 0.06 |
| | | $\mu_x = 6\mu_\lambda$ centroid | 4.2 | 0.1 | 4.95 | 0.05 |
| | | $\mu_x = 6\mu_\lambda$ random | 3.9 | 0.1 | 5.25 | 0.05 |
| | | $\mu_x = 12\mu_\lambda$ average | 3.6 | 0.2 | 7.10 | 0.10 |
| | | $\mu_x = 12\mu_\lambda$ centroid | 3.6 | 0.2 | 7.10 | 0.10 |
| | | $\mu_x = 12\mu_\lambda$ random | 3.4 | 0.2 | 7.40 | 0.10 |
| S5D | $\ln \mathrm{CV}_\lambda = a \ln L_\mathrm{s} + b$ | $k = D$ | 0.520 | 0.004 | −5.38 | 0.01 |
| | | $k = p, d, D$ | 0.510 | 0.006 | −5.01 | 0.02 |
| S5E | $\ln \mathrm{CV}_0 = a \ln L_\mathrm{s} + b$ | $k = d$ | 0.42 | 0.01 | −3.48 | 0.03 |
| | | $k = D$ | 0.53 | 0.01 | −4.52 | 0.05 |
| | | $k = p, d, D$ | 0.43 | 0.02 | −3.08 | 0.07 |
